# Nucleoid compaction during antibiotic stress excludes the SOS regulator LexA

**DOI:** 10.64898/2026.09.01.748551

**Authors:** Pilar Lörzing, Daria Miasoedova, Michael Schlierf

## Abstract

The bacterial SOS response promotes DNA repair, survival, and mutagenesis under genotoxic stress, especially during sub-lethal antibiotic exposure. This response is regulated by LexA, a transcriptional repressor controlling SOS gene expression, but how LexA coordinates this network as antibiotic stress alters nucleoid structure is unclear. Using ciprofloxacin-induced DNA damage, we investigated how increasing antibiotic stress affects the spatial relationship between LexA and the nucleoid in single *E. coli* cells. Through 3D single-molecule imaging and functional assays, we found that sublethal ciprofloxacin doses activated SOS, expanded nucleoids, and maintained LexA association. In contrast, higher stress caused severe DNA damage, compacted nucleoids, and LexA exclusion, yet some cells remained viable and recovered after drug removal. These results demonstrate that the SOS response involves both temporal and spatial regulation, with LexA and nucleoid organization adapting to damage severity to modulate SOS functions.

## Introduction

Bacteria must continuously adjust their physiology to survive in dynamic environments^1–3^. A major threat to cellular integrity arises from damage to the chromosome, which can originate from both endogenous processes and external stressors^4–7^. To counteract such challenges, bacteria employ dedicated regulatory networks that coordinate gene expression across the chromosome^8,9^. One of the best studied examples is the SOS response, a central transcriptional program activated by DNA damage^10,11^. The SOS response is controlled by the transcriptional repressor LexA. In the absence of DNA damage, LexA represses transcription by binding to operator sequences of more than 50 SOS genes distributed across the chromosome in *E. coli*, including factors involved in DNA repair, recombination, and cell-cycle control^12–15^ (Fig. 1). Upon genotoxic stress, the accumulation of single-stranded DNA promotes RecA filament formation, which stimulates LexA autocleavage and, by extension, relieves repression of the SOS regulon^16,17^. Notably, only unbound LexA can undergo cleavage, while DNA-bound LexA must first dissociate before it can be cleaved^18,19^. As LexA levels decline, individual promoters were shown to respond with distinct kinetics depending on their unbinding properties, creating a temporal order of gene induction^20^. Thus, e.g., high-fidelity repair pathways are activated first, followed by error-prone processes that promote survival under persistent stress^10,20,21^. Re-synthesis of LexA subsequently restores repression once DNA damage is resolved^22^. While the molecular mechanisms of SOS regulation are well characterized, an additional layer arises from the fact that LexA must coordinate repression of targets that are distributed across the chromosome. This raises the question of how regulatory function is linked to the spatial organization of the chromosome inside the bacterial cell (Fig. 1). The nucleoid represents a compact, yet, dynamic arrangement of the chromosome, composed of densely packed DNA together with associated proteins and RNA, that allows DNA-associated processes such as replication, transcription, and repair to occur within a confined cellular volume^23–27^. This organization is achieved through hierarchical compaction mechanisms, including DNA supercoiling, nucleoid-associated proteins, and higher-order folding into topological domains and macrodomains^28–31^. Despite this dense packing, the genome remains accessible for these DNA-associated processes and their regulation^27^. Importantly, nucleoid structure is not static but was shown to continuously remodel during growth and in response to stress^24,32,33^. Perturbations of DNA-associated processes, including those induced by antibiotics, lead to rapid changes in nucleoid morphology^34–37^. Because regulation of the SOS response requires coordination across multiple gene loci distributed throughout the chromosome, it is inherently linked to the spatial organization of the nucleoid. LexA must locate and bind its operator sites within the nucleoid, competing with other DNA-binding proteins while maintaining rapid response kinetics^13,38,39^. Single-molecule studies have shown that LexA occupies multiple functional states in the cell, including DNA-bound, transiently associated, freely diffusing, and cleaved forms, reflecting its roles in target search, repression, and turnover^40,41^. Since transcriptional repression must be coordinated across promoters distributed throughout the chromosome, the positioning and accessibility of LexA may constitute an additional regulatory layer beyond its biochemical properties. Changes in nucleoid architecture during DNA damage may therefore influence LexA positioning, promoter accessibility, and the ability of cells to efficiently induce SOS response. Yet, the spatial distribution of LexA within the nucleoid remains unexplored.

**Figure 1.**
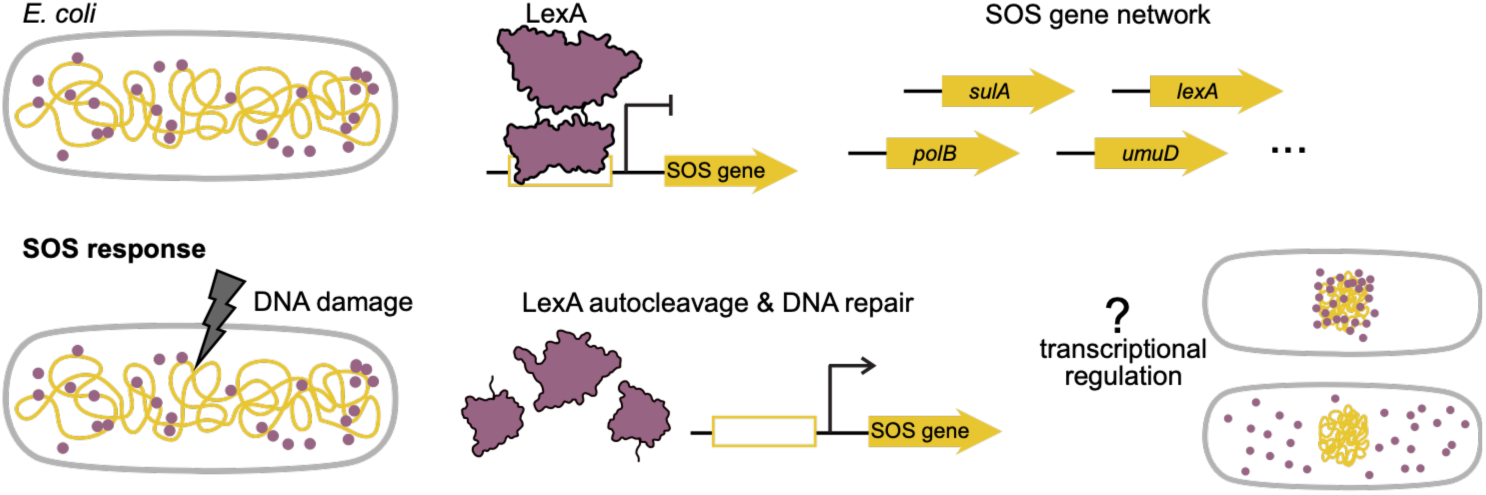
Organization of LexA in the bacterial SOS response. The SOS response coordinates DNA repair, mutagenesis, and survival following genotoxic stress. Under basal conditions, LexA (magenta) is distributed throughout the cell and represses SOS genes (yellow) across the chromosome by binding operator sequences. DNA damage induces LexA autocleavage, relieving repression of the SOS regulon. Because SOS promoters are distributed across the chromosome, LexA positioning may influence how repression is coordinated during the damage response.

Ciprofloxacin, a fluoroquinolone antibiotic that targets DNA gyrase in *E. coli*, generates DNA lesions that activate RecA-dependent LexA cleavage and thereby induce the SOS response^42^. Here, we use ciprofloxacin-induced DNA damage across a defined dose range, from sublethal exposure to strong antibiotic stress, to investigate how different levels of genotoxic stress reshape the spatial relationship between LexA and the nucleoid in single cells. By combining 3D single-molecule imaging of LexA with nucleoid morphology and measurements of SOS activity and survival, we link chromosome reorganization to the spatial coordination of SOS regulation across distinct damage states. We find that ciprofloxacin-induced SOS activation is accompanied by dose-dependent LexA–nucleoid reorganization, ranging from expanded nucleoids with partial LexA association at low ciprofloxacin exposure to compacted nucleoids with reduced LexA overlap under stronger antibiotic stress, while cells remain transcriptionally active and viable across the tested conditions.

## Results

To investigate how LexA positioning contributes to coordinating the SOS response within the spatially organized chromosome, we imaged the three-dimensional organization of LexA in *E. coli* and its relationship to nucleoid structure. We used a chromosomally-encoded LexA-PAmCherry (LexA^PA^) strain using λ-red recombineering^41^. We confirmed that the strain showed a wild-type like growth, unchanged ciprofloxacin susceptibility (Supplementary Fig. 1) and in *in vitro* assays LexA^PA^ showed identical operator affinity and no delay in RecA-mediated cleavage activity (Supplementary Fig. 3, 4). First, we imaged chromosomally expressed LexA^PA^ in fixed *E. coli* cells using 3D PALM^43^. Subsequently, we imaged the same cells with PAINT-based visualization of either the nucleoid (JF646-Hoechst) or the cell membrane^44^ (Potomac Red) (Fig. 2a, f). Reconstructed 3D PALM images revealed LexA^PA^ clusters within the cell (Fig. 2a). To exclude that this pattern resulted from fluorescent protein-dependent clustering, we also imaged LexA-Dendra2, a strictly monomeric fluorescent protein fusion and not associated with aggregation^45,46^, and cytosolic PAmCherry control. LexA-Dendra2 showed a similar clustered distribution, whereas PAmCherry was homogenously distributed throughout the cytosol (Supplementary Fig. 2). Thus, the observed pattern reflects likely LexA organization rather than aggregation of a fluorescent protein tag.

**Figure 2.**
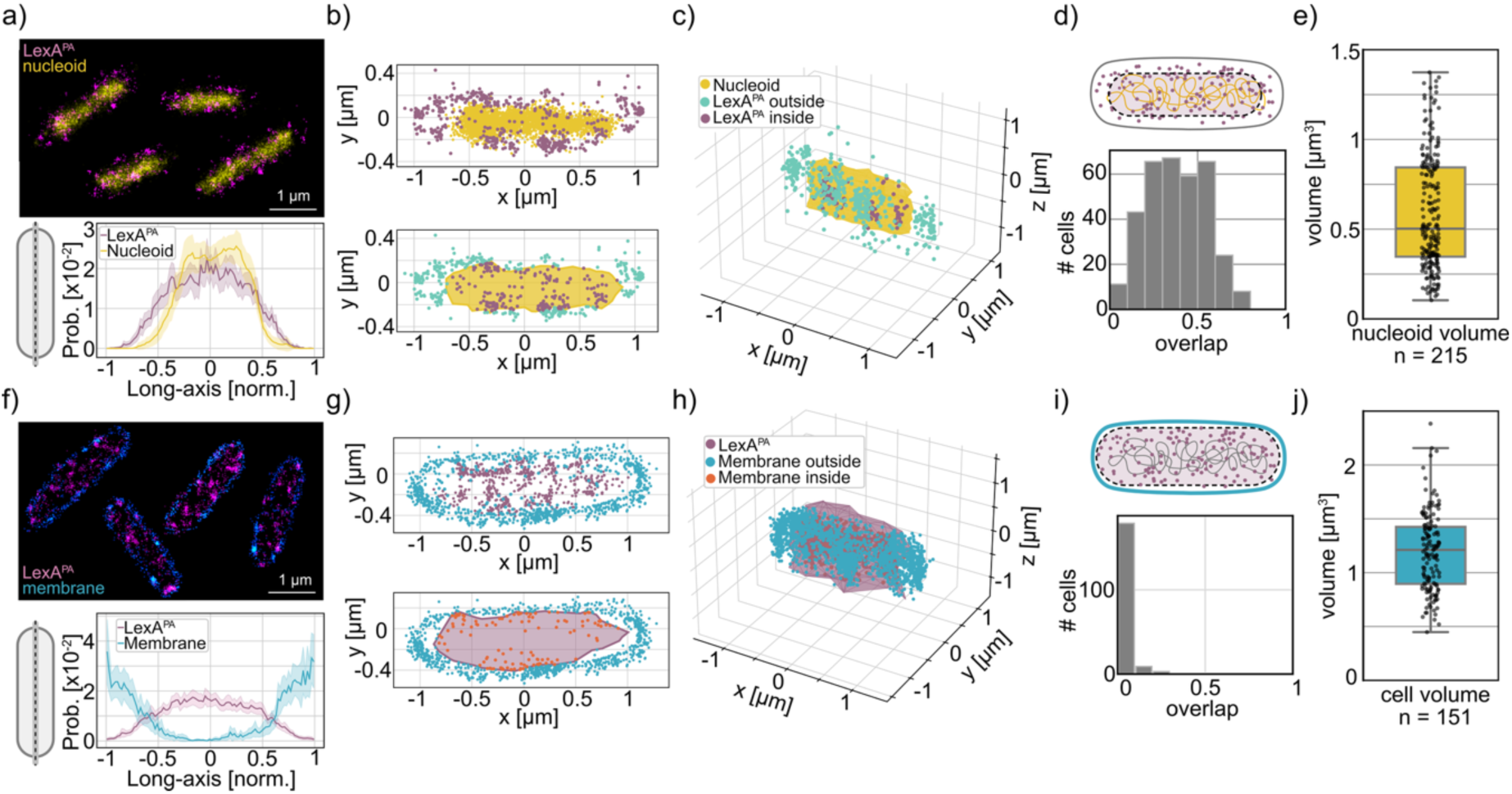
LexA^PA^ largely overlaps with nucleoid in unstressed cells. **(a)** Top: Dual-colour super-resolution images of LexA^PA^ (magenta) and the nucleoid labelled with JF646-Hoechst (yellow) in individual E. coli cells. Bottom: Mean normalized one-dimensional localization profiles of LexA^PA^ and the nucleoid along the cell long axis (n = 177; ±95% CI). **(b)** Two-dimensional colocalization of LexA^PA^ and the nucleoid (JF646-Hoechst). Top: representative cell shown as a scatter plot. Bottom: quantification using a concave hull constructed around nucleoid localizations; LexA^PA^ localizations are classified as overlapping (purple) or non-overlapping (turquoise) with the nucleoid region (yellow). **(c)** Three-dimensional colocalization analysis performed analogously to (a) using full 3D coordinates. **(d)** Global 3D colocalization, defined as the fraction of LexA^PA^ localizations within the nucleoid hull (mean ± SD). **(e)** Nucleoid volume per cell determined from concave hull reconstruction. **(f)** Top: Dual-colour super-resolution images of LexA^PA^ (magenta) and membrane staining with Potomac Red (blue). Bottom: Mean normalized 1D localization profiles of LexA^PA^ and the membrane (n = 174; ±95% CI) **(g)** 2D colocalization example of LexA^PA^ and the membrane (Potomac Red). Top: representative scatter plot. Bottom: quantification using a concave hull around LexA^PA^ localizations; membrane localizations are classified as overlapping (purple) or non-overlapping (turquoise) with the LexA^PA^-defined region (magenta). **(h)** 3D colocalization analysis performed analogously to (e). **(i)** Global 3D colocalization, defined as the fraction of membrane localizations within the LexA^PA^ hull (mean ± SD). **(j)** Cell volume per cell determined from concave hull reconstruction of membrane localizations.

LexA^PA^ was most abundant in the central, nucleoid-containing region, while also extending into the surrounding cytoplasm (Fig. 2a). To quantify the spatial relationship between LexA^PA^ and the nucleoid directly in three dimensions, we established a coordinate-based colocalization analysis. The reconstructed coordinate clouds can be used directly for geometric spatial analysis rather than relying on intensity-based overlap. For each cell, JF646-Hoechst localizations were used to reconstruct the nucleoid boundary by generating a concave hull around the localizations. As illustrated in two dimensions, LexA^PA^ localizations were assigned as either inside or outside the nucleoid hull (Fig. 2b). We then extended the same principle to the full 3D data set (Fig. 2c), generating a three-dimensional concave hull of the nucleoid and calculating the fraction of LexA^PA^ localizations contained within this volume. This analysis yielded on average a LexA^PA^–nucleoid overlap fraction of 0.38 ± 0.17 per cell (Fig. 2d). Importantly, the 3D nucleoid hull also yielded a nucleoid volume measure, giving a mean volume of 0.58 ± 0.32 µm^3^ per cell (Fig. 2e).

To validate the coordinate-based analysis against an independent cellular reference, we applied the same workflow to LexA^PA^ and the membrane PAINT label Potomac Red (Fig. 2f). Here, concave hulls were generated around the LexA^PA^ localizations, and Potomac Red localizations were classified as inside or outside these hulls. As expected, LexA^PA^ occupied the intracellular space with marginal overlap fraction with the membrane signal of 0.04 ± 0.04 (Fig. 2g, h, i). Simultaneously generated concave hulls from Potomac Red localizations delineated the cell boundary and provided a measure of cell volume (1.2 ± 0.36 µm^3^; Fig. 2j).

Having established this coordinate-based analysis in unstressed cells, we used it to examine LexA^PA^ spatial organization during SOS induction. Exponentially growing bacterial cells were treated with ciprofloxacin (CIP), a fluoroquinolone antibiotic causing DNA damage by inhibiting DNA gyrase and topoisomerase IV^42^ and known to induce the SOS response also at sub-lethal concentrations^47^. We determined the minimal inhibitory concentration (MIC) under our growth conditions to be 30 ng/ml (Supplementary Fig. 1c, d). *E. coli* cells were then exposed to 0.2x, 1x, 2x, and 5x MIC^CIP^ and fixed after 10, 30, 60, and 120 min of continuous treatment. Using dual color 3D SMLM, we monitored LexA^PA^ distribution and nucleoid morphology over time and across increasing levels of genotoxic stress (Fig. 3a). After 120 min of CIP treatment, reconstructed images showed elongated, filamentous cells at all tested concentrations (Fig. 3a, Supplementary Fig. 6), consistent with SOS-associated inhibition of cell division^48^. Using the membrane localization we quantified an approximate 1.5-fold increase in cell volume over 120 min (Fig. 3d).

**Figure 3.**
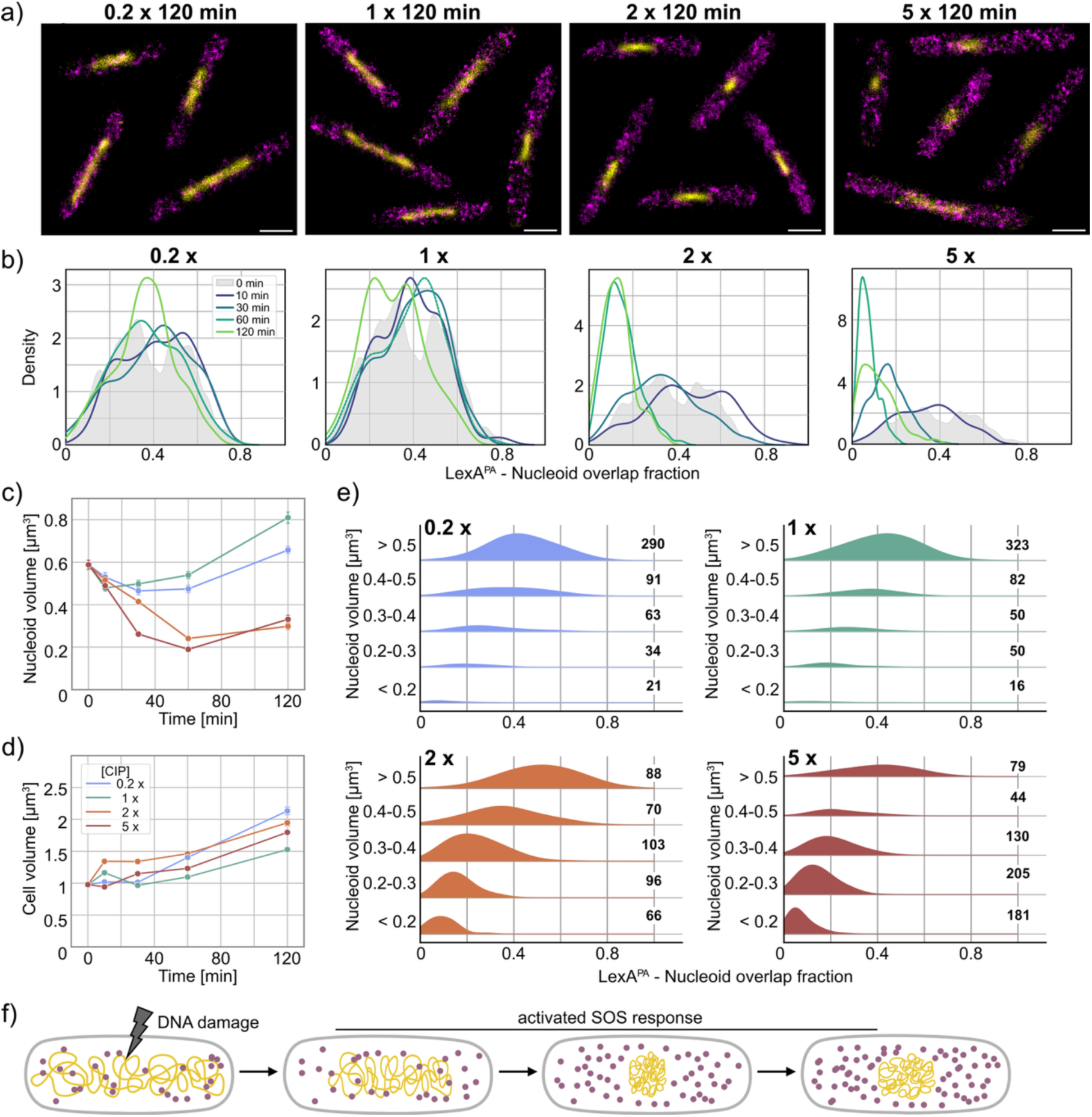
LexA^PA^ -nucleoid overlap scales with ciprofloxacin induced nucleoid compaction. **(a)** SOS response time course following treatment with CIP at 0.2x, 1x, 2x, and 5x MIC^CIP^ for 120 min. Shown are representative dual-colour super-resolution images of LexA^PA^ (magenta) and the nucleoid (JF646-Hoechst, yellow) in individual E. coli cells. Scale bar, 1 µm. **(b)** Time-dependent distributions of LexA^PA^– nucleoid overlap across ciprofloxacin concentrations. Kernel density estimates show the fraction of LexA^PA^ localizations within the 3D JF646-Hoechst nucleoid volume per cell for the indicated ciprofloxacin concentrations. Each panel represents one concentration. The 0 min reference distribution is shown in grey, and colored curves indicate subsequent treatment time points. **(c)** Mean nucleoid volume per cell derived from concave hull reconstruction of JF646-Hoechst localizations across the full time course (error bars, SE). **(d)** Mean cell volume per cell derived from concave hull reconstruction of Potomac Red-labelled membrane across the full time course (error bars, SE). **(e)** Ridgeline plots show distributions of the LexA^PA^ overlap fraction within the 3D JF646-Hoechst nucleoid volume per cell, grouped by nucleoid volume bins. Each panel shows one ciprofloxacin concentration. Ridge height is scaled by the number of cells per bin, with sample sizes indicated. **(f)** The schematic summarizes LexA redistribution during ciprofloxacin-induced SOS activation. Under basal conditions, LexA (magenta) is distributed throughout the cell and represses SOS genes across the chromosome (yellow). Strong ciprofloxacin stress leads to nucleoid compaction, increased LexA production, and LexA expulsion from the nucleoid.

We next examined how nucleoid morphology changed along the time course at different [CIP] and observed distinct nucleoid morphologies depending on the degree of genotoxic stress. At 0.2x and 1x MIC^CIP^, nucleoid volume initially remained close to untreated level during the first 60 min, before increasing at 120 min to 0.66 ± 0.24 µm^3^ and 0.81 ± 0.34 µm^3^, respectively (Fig. 3c). In contrast, higher CIP concentrations caused a pronounced compaction phenotype. At 2x and 5x MIC^CIP^, nucleoids formed dense structures near midcell (Fig. 3a), and compaction occurred earlier at the higher CIP dose (Fig. 3c, Supplementary Fig. 6). At 5x MIC^CIP^, mean nucleoid volume decreased to half its original volume in 30 min and reached a third of the original volume after 60 min, followed by a modest increase at 120 min, yet, remaining below the untreated level.

In parallel, we determined LexA^PA^ colocalization with the same time and dose dependence (Fig. 3b, Supplementary Fig. 7). During low-dose CIP exposure, LexA^PA^ remained associated with the nucleoid throughout the time course. At 0.2x MIC^CIP^, LexA^PA^-nucleoid overlap did not differ significantly from unstressed cells at any timepoint and was 0.36 ± 0.14 after 120 min. At 1x MIC^CIP^, overlap also remained comparable to the unstressed condition up to 60 min, but decreased significantly after 120 min to 0.31 ± 0.14. In contrast, higher CIP concentrations caused a pronounced displacement of LexA^PA^ from the nucleoid region. At 2x MIC^CIP^, overlap decreased significantly after 60 min to 0.15 ± 0.08 and remained low at 120 min (0.13 ± 0.07). At 5x MIC^CIP^, this transition occurred earlier, with a significant decrease to 0.17 ± 0.08 after 30 min, followed by 0.07 ± 0.04 at 60 min, and 0.12 ± 0.08 after 120 min (Fig. 3b, Supplementary table 1). In contrast, LexA^PA^ membrane overlap remained low throughout the CIP time course (Supplementary Fig. 8).

The population averages suggested that LexA^PA^ redistribution is linked to nucleoid morphology. We therefore analysed the relationship between nucleoid volume and LexA^PA^– nucleoid overlap on the single-cell level. Cells from each CIP condition were binned according to nucleoid volume, and the overlap distributions were compared within these bins (Fig. 3e). We found cells with expanded nucleoids showed high LexA^PA^ overlap, whereas cells with compacted nucleoids showed strongly reduced overlap. In low-dose conditions, the majority of cells populated large-volume nucleoids and maintained overlap values around 0.4. In high-dose conditions, cells were enriched in small-volume nucleoids, where overlap decreased to approximately 0.1. Concluding, low CIP stress produces an expanded nucleoid state in which LexA remains broadly associated with the chromosome. Higher CIP stress drives nucleoid compaction and a concomitant decrease of LexA^PA^ overlap with the compacted nucleoid (Fig. 3f).

To determine whether cells remained viable after CIP-induced nucleoid reorganization, we measured growth recovery and survival following antibiotic treatment (Fig. 4a). Cultures exposed to 0.2x-5x MIC^CIP^ for 10-120 min were washed and resuspended in fresh medium and growth curves were monitored. Increasing CIP concentrations resulted in progressively flatter growth curves and lower final OD_600_ values, as illustrated for a recovery after 30 min CIP exposure (Fig. 4b). Because bacterial biomass increases exponentially during log-phase growth, OD_600_ becomes linear after log transformation and the slope of this relationship provides the specific growth rate^49^. To quantify recovery across all CIP concentrations and exposure times, we determined the slope of the log-linear OD_600_ trace during the first 5 h after antibiotic removal, yielding an initial recovery rate (Fig. 4c). This fixed early window captures delayed growth resumption and subsequent outgrowth before cultures approach stationary phase. All conditions resumed growth after CIP removal, including cells exposed to 5x MIC^CIP^ for 120 min (Fig. 4b). Recovery, however, became progressively delayed and slower with increasing CIP concentration and treatment duration, as hypothesized. At 0.2x and 1x MIC^CIP^ growth rates remained only moderately reduced, with a significant reduction only after 120 min. In contrast, 2x and 5x MIC^CIP^ caused earlier and stronger reductions in recovery growth rates, becoming significant after 30 min and most pronounced after 120 min (Fig. 4c, Supplementary table 2). In parallel, we determined the number of colony forming units (CFU) corroborating that the reduction in recovery was accompanied by a dose- and time-dependent loss of viability (Fig. 4d). Survival at 0.2x MIC^CIP^ remained close to the untreated control. At 1x and 2x MIC^CIP^, CFU counts decreased gradually over time, with a significant reduction after 120 min. The strongest loss was observed at 5x MIC^CIP^ with a significant reduction already after 10 min and an approximately 300-fold reduction after 120 min. Nevertheless, colonies were recovered from all conditions (Supplementary table 3), showing that a viable fraction persisted across the CIP dose range.

**Figure 4.**
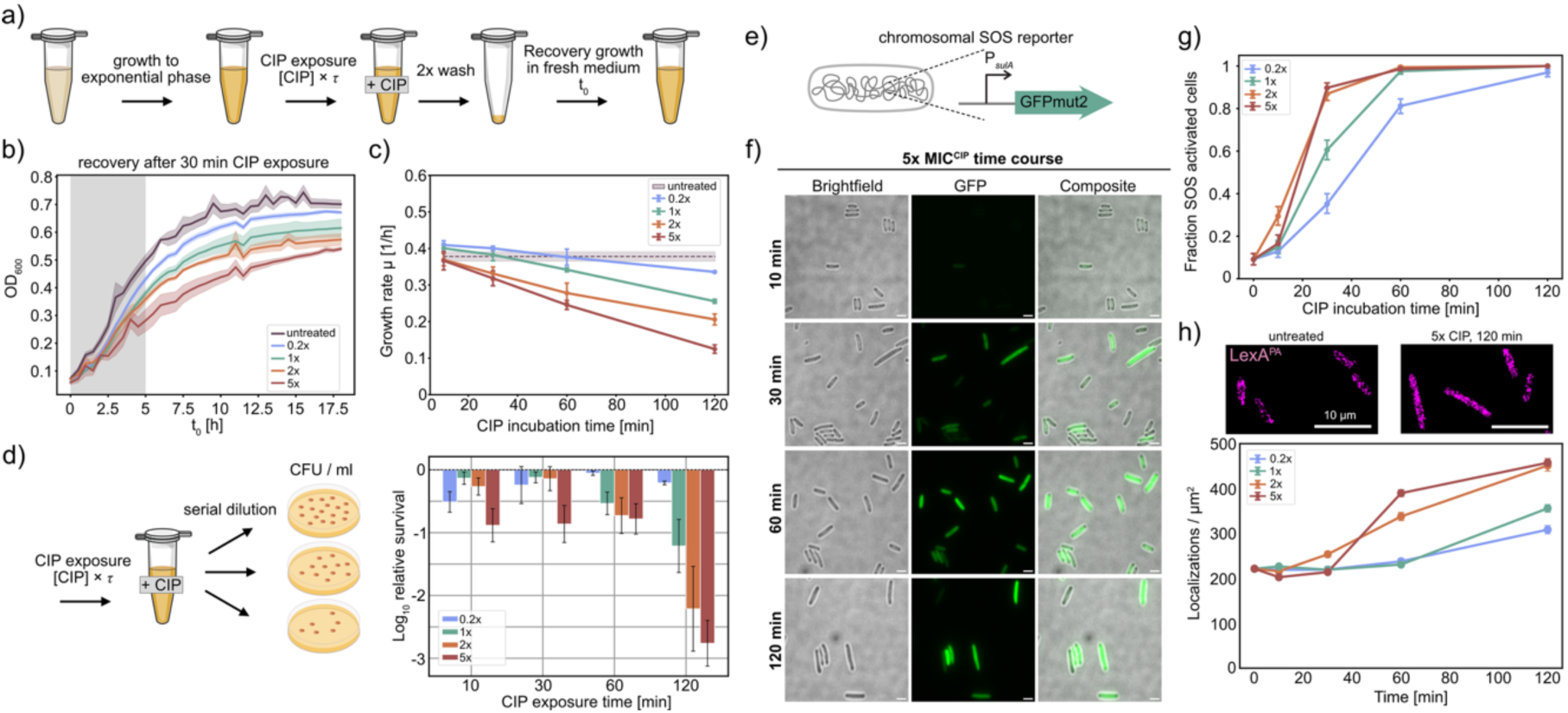
Cells remain transcriptionally active and viable across ciprofloxacin-induced DNA damage conditions. **(a)** Schematic of the recovery assay. Early exponential-phase E. coli cultures were treated with the indicated CIP concentrations for different durations, washed twice, resuspended in fresh medium, and monitored for regrowth by OD_600_ for 18 h. **(b)** Representative recovery growth curves after 30 min CIP exposure. Lines show the mean OD_600_ from biological triplicates; shaded areas indicate SD. **(c)** Growth rates determined from the initial recovery phase from t_0_[0–5 h] after CIP treatment. Points show mean growth rates for each concentration and exposure time; error bars indicate SD from biological triplicates. The dashed line indicates the untreated control. **(d)** Survival assay schematic and quantification. Early exponential-phase cultures were treated with CIP, serially diluted, plated on LB agar, and CFU/ml were determined after overnight incubation. Survival was calculated by normalizing CFU/ml to the mean untreated control at the corresponding time point. Bars show mean log_10_ survival fractions across biological replicates; error bars indicate SEM. **(e)** Schematic of the chromosomal SOS reporter strain expressing GFPmut2 under control of the sulA operator region. **(f)** Representative brightfield, GFP, and composite images of the SOS reporter strain during the 5x MIC^CIP^ time course. **(g)** Fraction of SOS-activated cells over time for the indicated CIP concentrations. Cells were classified as SOS activated when their mean sulA-GFP signal exceeded the cellular background mean by more than 3 SD. Points show activated fractions; error bars indicate bootstrapped 95% CIs. **(h)** Representative PALM reconstructions of LexA^PA^ in untreated cells and after 120 min 5x MIC^CIP^ treatment, with quantification of LexA^PA^ localization density across all conditions. Points show mean localizations per µm² cell area; error bars indicate bootstrapped 95% CIs.

To monitor SOS-dependent transcription throughout the CIP time course, we generated a chromosomally integrated P*_sulA_*-GFPmut2 reporter. GFPmut2 was previously characterized as a fast-maturing expression reporter, allowing us to cover also fast responses of the SOS response^50^. The reporter was constructed by λ Red recombineering and integrated as a single copy at the attλ site of the *E. coli* chromosome (Fig. 4e). In this construct, GFPmut2 expression is driven by the LexA controlled *sulA* promoter, while the native *sulA* gene remains unmodified. GFP signal therefore reports loss of repression of the *sulA* promoter following DNA damage^4,51,52^. In turn, we could determine whether cells retained SOS transcriptional activity at the differently compacted nucleoid states. Subsequently, cells were treated with CIP under identical conditions as previously, and the mean GFP intensity was quantified per cell (Supplementary Fig. 10). We then classified a fraction of SOS-activated cells when the GFP signal crossed a threshold obtained from the non-reporter control (Fig. 4f). Notably, an increase in the GFP signal was detectable in a subset of cells already after 10 min of CIP treatment and grew with both exposure time and CIP concentration (Fig. 4g). At 2x and 5x MIC^CIP^, the SOS-active fraction of cells increased rapidly with nearly the entire population becoming SOS active by 60 min. At 0.2x MIC^CIP^, induction was slower but increased continuously over the time course, reaching a high fraction of SOS-active cells after 120 min. Since the *lexA* gene itself is SOS regulated^10,53^, the LexA^PA^ density provides a complementary measure of the SOS associated expression. Thus, we determined independently the LexA^PA^ localization density for the same time courses (Fig. 4g). Despite the pronounced cell elongation during CIP treatment, LexA^PA^ density increased over time. At 2x and 5x MIC^CIP^, density increased strongly from 30 min onwards and doubled after 120 min. In contrast, LexA^PA^ density rose more gradually at 0.2x and 1x MIC^CIP^.

## Discussion

Because LexA represses SOS promoters distributed across the chromosome, SOS regulation must operate within a nucleoid that is itself reorganized by DNA damage. Here, we combined 3D single-molecule imaging of LexA^PA^ and the nucleoid and membrane with measurements of SOS reporter activity, recovery, and survival to relate spatial organization to the response to ciprofloxacin-induced damage. In untreated cells, LexA^PA^ was distributed throughout the intracellular space but ∼ 40% remained associated with the nucleoid, likely either bound to its operators or in search for an empty operator. Previous single-molecule studies identified both DNA-bound and freely diffusing LexA populations in unstressed cells^40,41^. Consistent with this, the partial LexA^PA^-nucleoid overlap observed here suggests that LexA is distributed between a nucleoid-associated fraction and a freely diffusing cytoplasmic pool. In addition to the spatial overlap, the 3D PAINT enabled single-cell quantification of nucleoid and cell volume. Untreated cells showed mean nucleoid and cell volumes of 0.58 ± 0.30 µm^3^ and 1.20 ± 0.36 µm^3^, respectively, consistent with reported values for exponentially growing *E. coli* ^54–56^. The broad distributions likely reflect the heterogeneous replication and cell-cycle states of the unsynchronized population (Supplementary Fig. 5).

Nucleoid morphology is shaped by chromosome-intrinsic factors, including supercoiling^24^ and DNA-binding proteins^29^, but also by the physical dimensions of the cell and the crowded cytoplasmic environment^27^. In bacteria, nucleoid size was shown to scale with cell size across different growth conditions, independently of DNA content^57^, and the available longitudinal space within the cell can further contribute to chromosome extension^58^. CIP exposure revealed distinct damage-dependent changes in nucleoid organization. At 0.2 x and 1x MIC^CIP^ exposure, nucleoid volume increased beyond 30 min while LexA^PA^ retained partial overlap with the nucleoid. This change coincided with an increase in cell volume. The enlarged cell volume may therefore contribute to the more extended nucleoid morphology observed at 0.2x MIC^CIP^. This state was accompanied by increased SOS reporter activity, showing that the SOS response was upregulated despite spatial proximity of LexA^PA^ to the nucleoid. Thus, the low-dose state may represent a condition in which SOS transcription is activated while the chromosome remains broadly accessible and LexA remains spatially associated with DNA.

In contrast, stronger CIP exposure resulted in a different response. At 2x and 5x MIC^CIP^, nucleoid volume decreased rapidly and LexA^PA^-nucleoid overlap was strongly reduced, yielding dense nucleoids from which LexA was excluded. Nucleoid compaction has been observed in *E. coli* following diverse genotoxic stresses, including UV irradiation, mitomycin C, bleomycin, and fluoroquinolone treatment and these compacted DNA states have been proposed to stabilize damaged chromosomes and support DNA repair^59–62^. In particular, Vikedal et al.^62^, showed that severe CIP-induced damage drives an ordered transition towards a dense midcell nucleoid. Their work identified RecN and RecA as essential components of this process and proposed that supercompaction forms part of an active response to double-strand-break damage. Our data add the SOS master regulator to this high-damage state: LexA was excluded from the compacted DNA, and therefore from SOS operator sites located within this region, while SOS reporter activity remained high. Thus, the high-CIP state is not only characterized by chromosome compaction, but also by strong loss of SOS gene repression, consistent with continued expression of repair and tolerance functions during severe DNA damage.

The CIP treatments that generated these LexA^PA^-nucleoid states still contained cells able to recover after drug removal: cultures from all conditions resumed growth, and colonies were recovered even after the strongest treatment. At 0.2x MIC^CIP^, SOS was induced with little loss of viability, showing that subinhibitory CIP triggers DNA-damage responses in a surviving population. With increasing CIP dose and exposure time, recovery growth and CFU counts declined. Thus, LexA-nucleoid states reflect different levels of bacterial survival under DNA damage rather than irreversible structural endpoints. This makes the low-dose state particularly relevant to fluoroquinolone resistance evolution. By inducing SOS without substantial killing, sub-MIC CIP exposes a surviving population to SOS-regulated repair and tolerance processes, some of which, including error-prone translesion synthesis, can be mutagenic. Their use may increase genetic variation and thereby the likelihood that resistant variants emerge under selection, without implying that SOS induction itself produces resistance^47,63^.

## Methods

### Culture Conditions

*E. coli* MG1655 chromosomally expressing LexA-PAmCherry fusion construct^64^ was grown in M9 minimal medium (1 x M9 salts [Sigma], 1mM MgSO4, 0.1 mM CaCl2, 0.4 % glucose) supplemented with 0.001 % biotin and thiamine. Cultures were grown in 1.5 ml tubes with a punctured lid at 37 °C until early exponential phase.

### Survival assay

Cell survival after ciprofloxacin treatment was quantified by colony-forming unit (CFU) analysis. *E. coli* MG1655 was grown overnight from a single colony in M9 minimal medium at 37 °C. The overnight culture was diluted 1:36 into fresh M9 medium and grown for 3.5 h to early exponential phase. Ciprofloxacin was added at the indicated concentrations, followed by incubation for the respective treatment times. Cultures were then serially diluted in M9 minimal medium, plated on LB agar, and incubated overnight at 37 °C. Colonies were counted the following day, and CFU/ml was calculated by correcting for the plated volume and dilution factor. Survival fractions were calculated for each ciprofloxacin-treated condition by normalizing CFU/ml values to the untreated control at the corresponding time point. Survival fractions were log10-transformed, and mean values with standard errors were calculated from biological replicates.

### Recovery assay

*E. coli* MG1655 was grown overnight from a single colony in M9 minimal medium at 37 °C. The overnight culture was diluted 1:36 into fresh M9 medium and grown for 3.5 h to early exponential phase. Ciprofloxacin was added at the indicated concentrations, and cultures were incubated for the respective exposure times. Cells were then washed twice in fresh M9 medium by centrifugation at 7000 × g for 1 min, removal of the supernatant, and resuspension of the pellet in fresh medium. After the second wash, cells were resuspended in 1 ml M9 medium, transferred to a 24-well plate, and recovery growth was monitored at 37 °C with shaking for 18 h using a plate reader (CLARIOstar^®^ Plus, BMG Labtech). OD_600_ was measured every 30 min. Recovery growth rates were calculated from the first 5 h of growth after transfer to fresh medium. For each replicate well, ln(OD_600_) was plotted against time and fitted by linear regression. The resulting slope was defined as the growth rate µ. Mean growth rates and standard deviations were calculated from biological triplicates. All stressed conditions were compared with the untreated control with multiple comparisons correction.

### Sample Preparation

Imaging was done in fixed cells. For fixation, bacteria were cooled on ice for 10 min and incubated with 4% paraformaldehyde in PBS, pH 7.4, for 35 min at room temperature, then washed twice with PBS, pH 7.4 and stored at 4 °C. Fixed *E. coli lexA-lk-PAmCherry* cells were permeabilized with 0.5% Triton X-100 in PBS, pH 7.4, to allow access for PAINT labels. Cells were pelleted by centrifugation at 7000 g for 1 min, resuspended in 400 µl of 0.5% Triton X-100, and incubated at room temperature for 45 min with shaking at 500 rpm.

Coverslips were cleaned by sonication in 5% Mucasol for 15 min, rinsed with milli Q water, and sonicated in HPLC-grade ethanol for 15 min. The washing step was repeated twice. Coverslips were blow dried with nitrogen gas and kept in a clean, dry glass slide holder until further use. Well chambers were prepared from 8-well sticky slides (Ibidi) using Poly-L-lysine (PLL) coated coverslips. Cleaned coverslips were incubated with 0.01% PLL for 30 min at room temperature, dried at 60 °C, aligned with the sticky slide, and pressed onto the adhesive side. Assembled chambers were incubated at 37 °C for >8 h for sealing.

Fixed and permeabilized cells were mixed with 100 nm Gold Nanoparticles (Alfa Aesar) as fiducial marker at a concentration of 1:20 v/v. The sample was added to each well and allowed to settle for 30 min at room temperature. Wells were washed twice with PBS to remove residual Triton X-100. PAINT label solution prepared in PBS, pH 7.4, was added for imaging. PAINT label concentrations were determined by initial titration experiments. Final concentrations were 300 pM for JF646-Hoechst and 450 pM for Potomac Red.

### Chromosomal SOS reporter strain construction

The pR6K-sulA-GFPmut2 plasmid was generated by Gibson assembly. The sulApΩgfp-mut2 insert was amplified from pZA31-sulApΩgfp-mut2^51^ using the primer pair *sulA_GFP_in_fwd* and *sulA_GFP_in_rev*, which introduced 30 bp homology regions to the pR6K backbone. The pR6K backbone was linearized using primers *pR6K_lin_fwd* and *pR6K_lin_rev*. Assembly was performed using the NEBuilder® HiFi DNA Assembly Reaction Protocol into *E. coli* DH5a cells.

Chromosomal insertion was carried out by λ Red recombineering using a protocol adapted from previously described methods^65,66^. For insertion at the attλ locus, pR6K-sulA-GFPmut2-lox71-cm-lox66 was linearized using primers *sulA_GFP_rec_fwd* and *sulA_GFP_rec_rev*, which introduced homology regions flanking the attλ site. *E. coli* MG1655 cells carrying pSC101-BAD-γβαA were induced with 0.25% L-arabinose to express the λ Red recombination genes. Competent cells were electroporated with 250 ng of the linearized PCR product. The temperature-sensitive helper plasmid was recovered by overnight incubation at 37 °C. Transformants were selected on LB agar supplemented with chloramphenicol (15 μg/ml) and successful chromosomal insertion was verified by colony PCR using primers *sulA_GFP_seq_fwd* and *sulA_GFP_seq_rev*. The chloramphenicol resistance cassette was subsequently removed by Cre-lox mediated recombination using the temperature-sensitive plasmid pSC101-BAD-Cre. Cre expression was induced with 5 mM L-arabinose. Temperature-sensitive plasmids were removed during incubation at 37 °C.

### Single molecule localization microscopy

3D SMLM experiments were carried out Nikon Eclipse Ti2-E inverted microscope (Nikon) system equipped with an ORBITAL-100 Ring TIRF system (Visitron Systems) and controlled with VisiView Software (Visitron Systems). A 100x Oil immersion objective (CFI TIRF Apochromat, NA 1.49, WD 0.12 mm, Nikon) was used with an additional 1.5x tube lens to increase the magnification. The detector was a Prime-95B Back-Illuminated scientific CMOS camera (Teledyne Photometrics). A cylindrical lens (0.67 m focal length, Cairn research) was inserted into the detection path to allow 3D astigmatic imaging.

PAmCherry was continuously activated with a 405 nm laser (0 – 1 mW, before objective) and excited at 561 nm with 8 mW (before objective) under HILO conditions. A time-lapse movie of 8,000 frames was recorded at a frame rate of 50 fps. Emission light was filtered by a 605/70 bandpass filter (AHF). PAINT labels were gifted from Lavis lab and Open Chemistry team (HHMI Janelia Research campus). Labels were prepared in 1x PBS (pH 7.4) from stock solution. Time-lapse movies of 8,000 – 9,000 frames were recorded with frame rates: JF646-Hoechst at 50 fps and Potomac Red at 33 fps. Both dyes were excited using 637 nm laser at 10 mW (before objective). Emission light was filtered using a dual-line laser rejection filter for 561 and 640 nm excitation light (ZET561/640, AHF). Brightfield images of the same field of view were taken before data acquisition.

### Raw SMLM data processing

Raw SMLM movies were processed using SMAP^67^. For sCMOS-specific single-molecule fitting, pixel-wise gain, offset, and noise maps were generated using the photon-free camera characterization workflow described by Diekmann et al.^68^, which were subsequently applied during single-molecule localization in SMAP. The shape of the 3D point spread function was calibrated from 10 z-stacks (10 nm step size) of 0.2 µm tetra speck fluorescent beads (Invitrogen). After localization, the data was corrected for drift using the drift correction plugin and the gold nanoparticles. Localizations across consecutive frames were correlated to accurately capture individual single molecules. A fixed maximum distance of 20 nm and up to 5 dark frames between localizations were used for merging to account for fluorophores emitting across multiple frames and possible blinking events.

### Channel alignment

PAINT images were registered to the PALM channel using a Fourier transform-based image registration approach, as described by Spahn et al.^69^. Translational shifts between channels were determined by frequency-domain cross-correlation, implemented in Python using the scipy.fft module for Fast Fourier Transforms. The calculated x- and y-shifts were applied to align the channels before further analysis. Cells were then segmented, and localizations were assigned to individual cells based on the segmented cell area, as described previously^43^.

### Colocalization analysis

Colocalization was determined using concave hulls (α-shapes) implemented in Python. For LexA-PAmCherry/JF646-Hoechst data, cells with fewer than 2000 Hoechst localizations were excluded. JF646-Hoechst localizations were reconstructed as 2D and 3D concave hulls using Delaunay triangulation with SciPy (scipy.spatial.Delaunay), with a 250 nm cutoff radius to exclude large empty regions. In 2D, triangles were combined into polygons using Shapely package; in 3D, connected tetrahedra were used to define the hull and calculate nucleoid volume. Global colocalization was quantified as the fraction of LexA-PAmCherry localizations inside the JF646-Hoechst hull. For LexA-PAmCherry/Potomac Red data, cells with fewer than 5000 Potomac Red localizations were excluded. The same analysis was performed using LexA-PAmCherry hulls, and colocalization was quantified as the fraction of Potomac Red localizations inside the LexA-PAmCherry hull. For 2D analysis, Potomac Red localizations were restricted to ±150 nm in z to retain the mid-cell plane. Cell volume was determined from 3D Potomac Red concave hulls.

### SOS GFP reporter analysis

SOS reporter activity was measured in fixed *E. coli* MG1655 cells carrying the chromosomal *sulApΩgfp-mut2* reporter. Cells were grown, treated with ciprofloxacin, and fixed as described above for the LexA-PAmCherry expressing strain. GFPmut2 fluorescence was imaged on the same microscope setup described above. For each field of view, a single GFP image was acquired using 488 nm excitation at 1 mW before the objective and an exposure time of 50 ms. Emission light was filtered using a quad-band filter set (ZET405/488/561/640m-TRFv2, Chroma). A corresponding brightfield image was acquired for each field of view and used for cell segmentation.

SOS induction was quantified from single-cell fluorescence measurements of the chromosomal sulA-GFP reporter imaging. Cells were segmented from brightfield images, and the resulting masks were used to measure the mean GFP intensity per cell in Fiji/ImageJ. Background fluorescence was estimated from a non-fluorescent wild-type control measured under identical conditions. For each condition, the mean and standard deviation of the *E. coli* MG1655 wildtype signal were calculated and used for background correction and thresholding. Single-cell GFP values were corrected by subtracting the mean background signal. Cells were classified as SOS-positive when their raw mean GFP intensity exceeded the background mean plus four standard deviations. The fraction of SOS-positive cells was calculated for each condition, and 95% confidence intervals were estimated by bootstrap resampling of the single-cell classifications with 1000 iterations.

### Statistics and Reproducibility

Statistical analyses were performed in Python. Survival and recovery data were analyzed using Dunnett’s test (scipy.stats.dunnett) against the respective untreated control. LexA^PA^-Hoechst colocalization was analyzed using Kruskal-Wallis tests (SciPy) followed by Dunn’s post hoc test with Holm correction (scikit-posthocs). Statistical significance was defined as p < 0.05. Biological replicates were used for survival and recovery assays; for microscopy analyses, n denotes individual cells. Sample sizes are indicated in the corresponding supplementary tables.

## Data Availability

The data supporting this study are available within the article and Supplementary Information. Additional datasets will be deposited in a public repository upon publication.

## Acknowledgements

We thank all members of the Schlierf lab for lively discussions during the development of this project. We thank Prof. Francis Stewart and Dr. Frank Groß (BIOTEC, TU Dresden) for the kind gift of the recombineering plasmids and Prof. Yixin Zhang (B CUBE, TU Dresden) for access to the plate reader. We thank Dr. Leonard Schärfen for the MG1655-LexA-PAmCherry strain. This project was partially supported by the Deutsche Forschungsgemeinschaft (DFG, German Research Foundation) under Germanýs Excellence Strategy – EXC-2068 – 390729961-Cluster of Excellence Physics of Life of TU Dresden and TU Dresden core funds (to M.S.).

## Author contributions

Conceptualization: MS, PL

Methodology: PL, DM

Investigation: PL, DM

Visualization: PL

Supervision: MS

Writing—original draft: PL

Writing—review & editing: PL, DM, MS

