## Supplemental Figures and Methods for "Nucleoid compaction during antibiotic stress excludes the SOS regulator LexA"

### Supplementary Figures

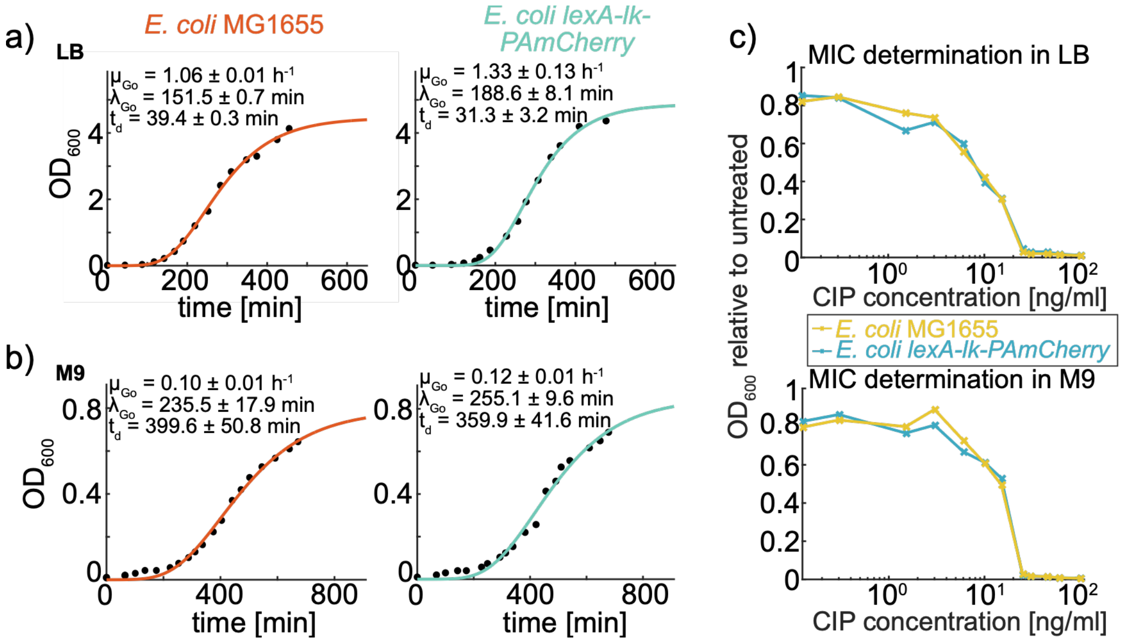

***Supplementary Figure 1 Growth curves and ciprofloxacin sensitivity of E. coli MG1655 and E. coli lexA-lk-PAmCherry.***

*(a) Growth curves in LB medium and (b) in M9 medium. Data were fitted with the Gompertz model to extract lag phase (λ), maximum specific growth rate (μ), and doubling time ($t_d$). (c) Minimal inhibitory concentration (MIC) of ciprofloxacin determined by broth dilution and $OD_{600}$*

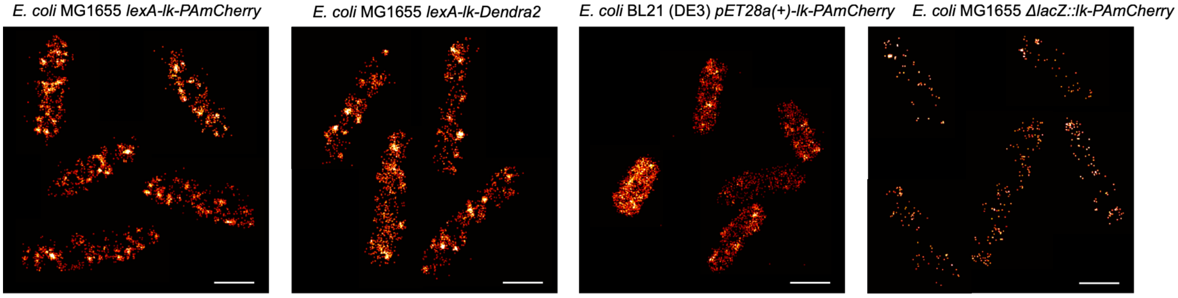

***Supplementary Figure 2 Super-resolution imaging of LexA fusion proteins and fluorescent controls.*** *Reconstructed super-resolution images of E. coli MG1655 LexA-lk-PAmCherry, E. coli MG1655 LexADendra2, E. coli MG1655 ΔlacZ::lk-PAmCherry, and E. coli BL21 (DE3) pET-28a(+)-lk-PAmCherry are shown for comparison. Scale bar: 1µm.*

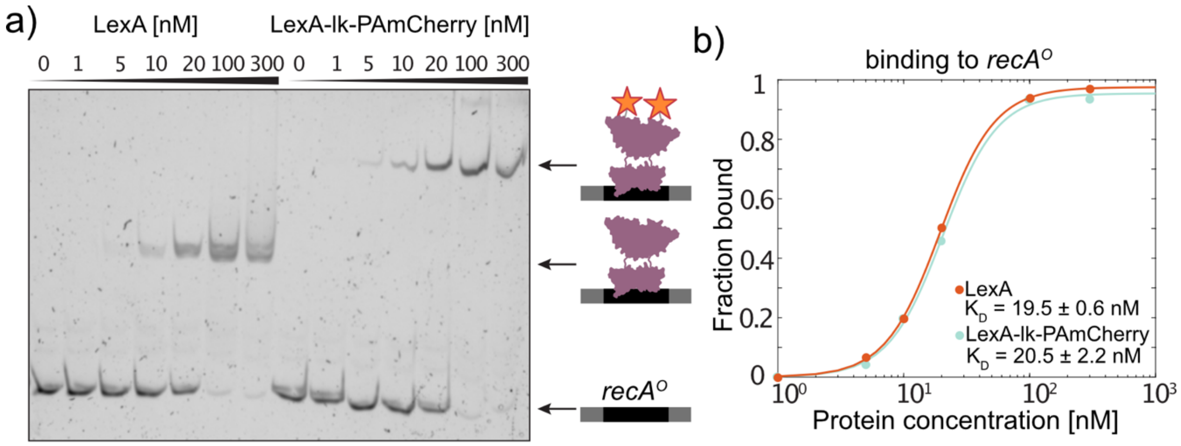

***Supplementary Figure 3 EMSA of LexA and LexA-PAmCherry binding to the recA operator.*** *(a) Electrophoretic mobility shift assay showing binding of LexA and LexA-lk-PAmCherry to recA operator DNA at the indicated increasing protein concentrations. Schematics indicate the positions of free DNA, LexA-bound DNA, and LexA-lk-PAmCherry bound DNA. (b) Quantification of EMSA band intensities. The fraction of bound DNA is plotted as a function of protein concentration, with fitted binding curves shown.*

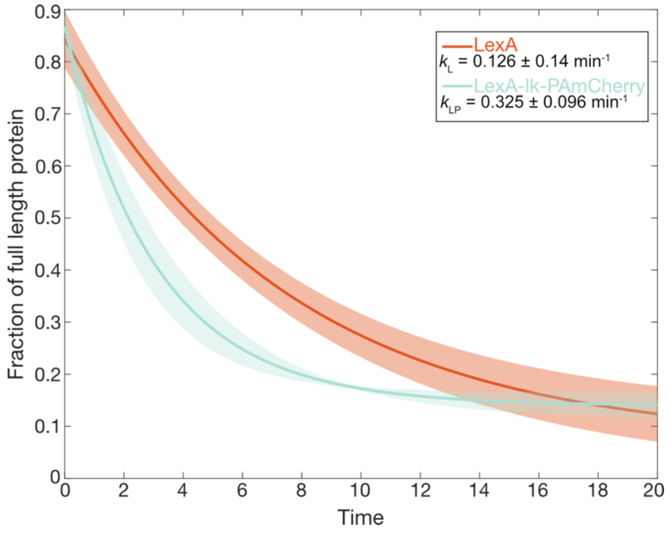

***Supplementary Figure 4 Quantification of LexA and LexA-PAmCherry cleavage.*** *Full-length LexA or LexA-lk-PAmCherry and the corresponding cleaved C-terminal fragments were used to determine the fraction of full-length protein during the cleavage reaction. Data represent the mean ± SD from three independent cleavage reactions. Mean values were fitted with a single-exponential decay function***,*** *where k represents the decay rate constant. Solid lines indicate the fitted mean and shaded areas indicate the SD.*

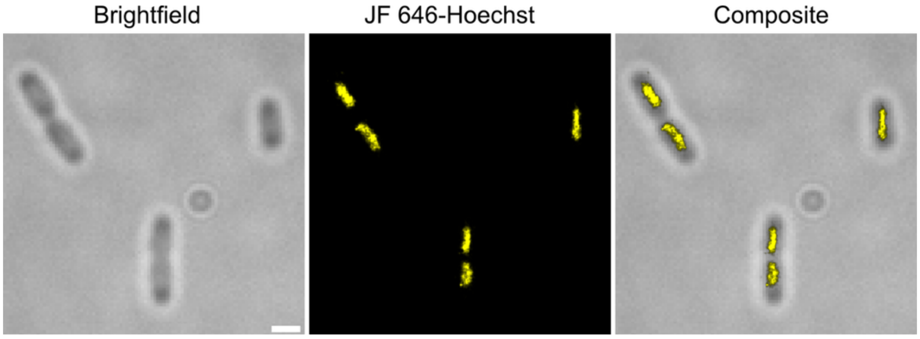

***Supplementary Figure 5 PAINT imaging of the bacterial nucleoid in individual E. coli cells.*** *Example field of view (FOV) from JF646-Hoechst imaging showing the brightfield image (left), reconstructed super-resolution image of the nucleoid (middle), and composite overlay (right). The FOV shows cell-to-cell variability in nucleoid morphology within the unsynchronized population, e.g. heterogeneous replication and cell-cycle states. Scale bar: 1µm.*

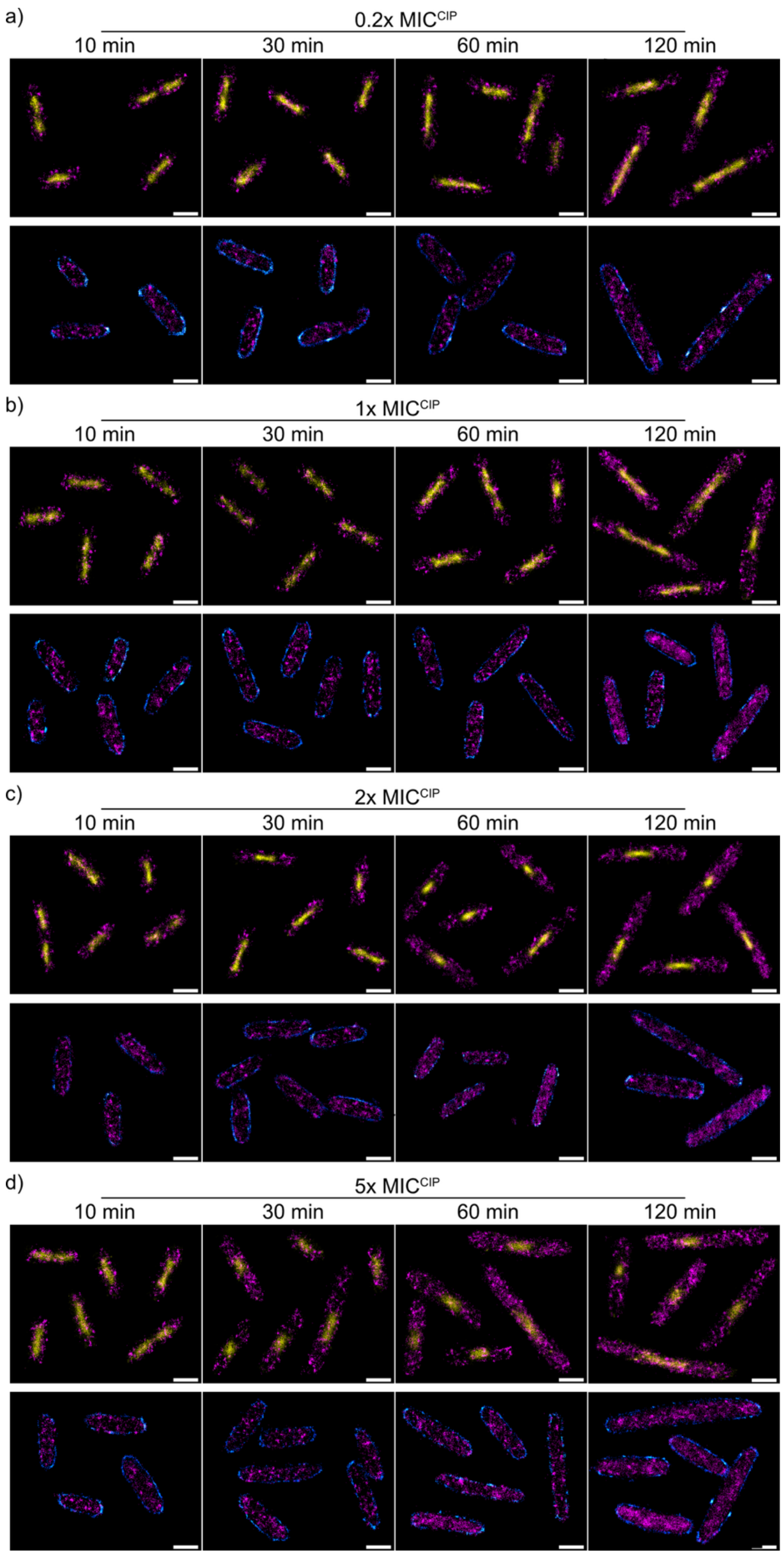

***Supplementary Figure 6 Time course dual-colour SMLM of LexA-lk-PAmCherry relative to the nucleoid and membrane during ciprofloxacin induced SOS response.*** *E. coli cultures were treated with ciprofloxacin at (a) 0.2x, (b) 1x, (c) 2x, and (d) 5x MIC$^{CIP}$ and fixed at defined time points (10, 30, 60, and 120 min). For each concentration, the upper panel shows reconstructed super-resolution images of LexA-lk-PAmCherry (magenta) and the JF646-Hoechst labelled nucleoid (yellow) at the indicated time points. Corresponding dual-colour images of LexA-lk-PAmCherry (magenta) relative to the Potomac Red labelled membrane (blue) are shown in the lower panel. Scale bar: 1 µm.*

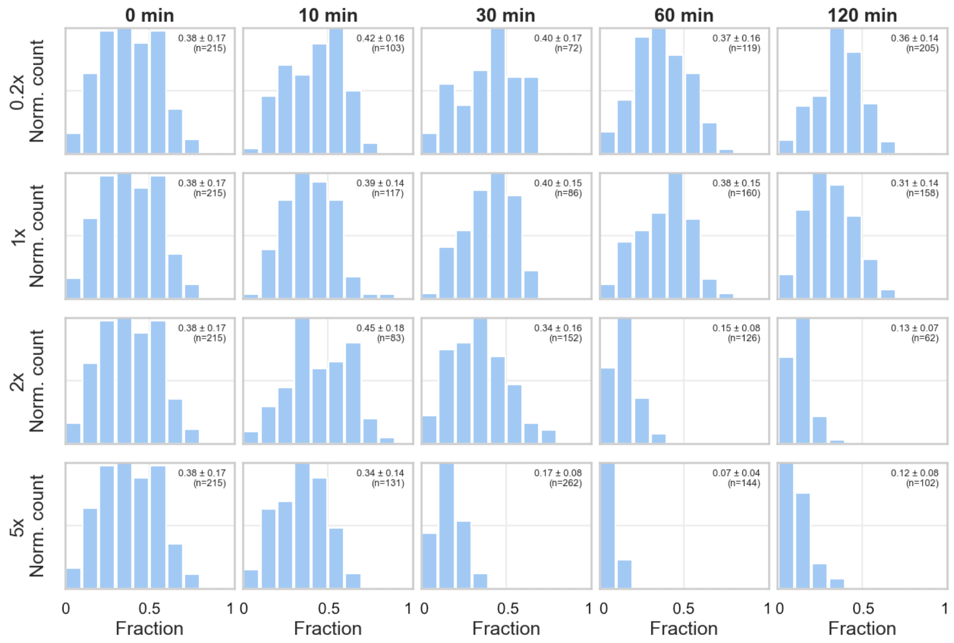

***Supplementary Figure 7 Three-dimensional colocalization of LexA-lk-PAmCherry with the nucleoid during the ciprofloxacin-induced SOS response.*** *Quantification from dual-colour SMLM imaging of LexA-lk-PAmCherry and JF646-Hoechst-labelled nucleoids Histograms show the distribution of the fraction of LexA-lk-PAmCherry localizations within the nucleoid area per cell for 0.2x, 1x, 2x, and 5x MIC$^{CIP}$ (top to bottom) at the indicated time points. The 0 min condition represents unstressed cells for comparison. Mean ± SD is indicated for each condition; n denotes the number of cells analyzed.*

***Supplementary Table 1*** *Pairwise statistical comparison of LexA-lk-PAmCherry - nucleoid colocalization between untreated cells (0 min) and each ciprofloxacin treatment timepoint. P-values were determined using Dunn's multiple-comparison test with Holm correction. Significance levels are indicated as ****, $p < 0.0001$; ***, $p < 0.001$; **, $p < 0.01$; *, $p < 0.05$; and ns, not significant.*

| **MIC$^{CIP}$** | **timepoint** | **Mean ± SD overlap** | **Significance** |
|---|---|---|---|
| 0x | 0 min | 0.381 ± 0.167 | - |
| 0.2x | 10 min | 0.417 ± 0.160 | ns |
| 0.2x | 30 min | 0.404 ± 0.170 | ns |
| 0.2x | 60 min | 0.369 ± 0.156 | ns |
| 0.2x | 120 min | 0.357 ± 0.138 | ns |
| 1x | 10 min | 0.388 ± 0.144 | ns |
| 1x | 30 min | 0.397 ± 0.145 | ns |
| 1x | 60 min | 0.382 ± 0.150 | ns |
| 1x | 120 min | 0.313 ± 0.136 | *** |
| 2x | 10 min | 0.450 ± 0.182 | ns |
| 2x | 30 min | 0.339 ± 0.160 | ns |
| 2x | 60 min | 0.149 ± 0.077 | **** |
| 2x | 120 min | 0.131 ± 0.072 | **** |
| 5x | 10 min | 0.344 ± 0.142 | ns |
| 5x | 30 min | 0.167 ± 0.079 | **** |
| 5x | 60 min | 0.071 ± 0.041 | **** |
| 5x | 120 min | 0.117 ± 0.080 | **** |

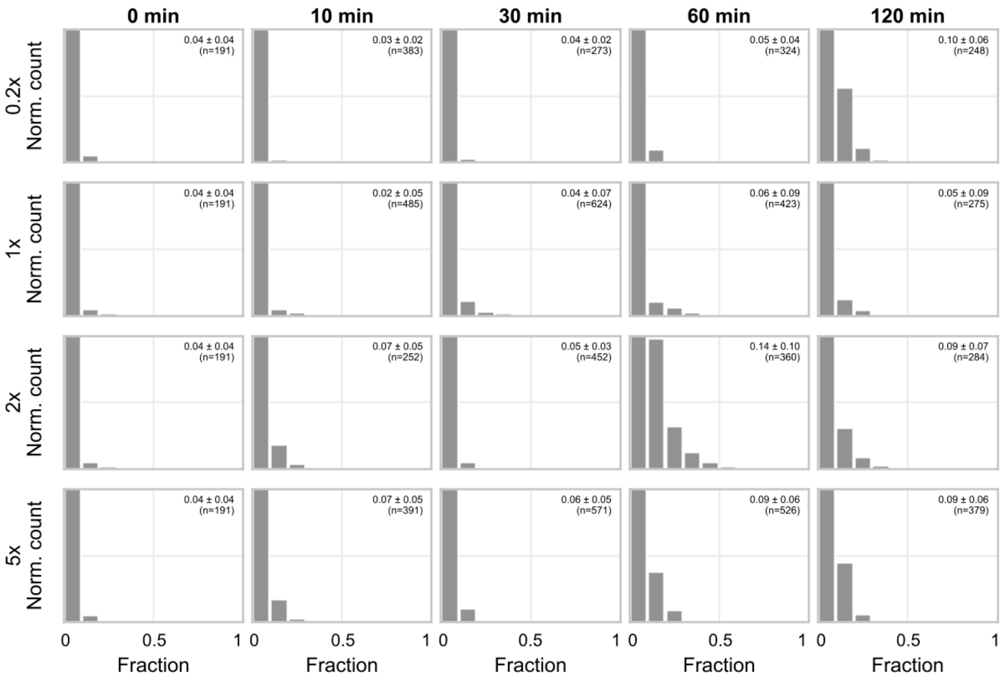

***Supplementary Figure 8 Three-dimensional colocalization of LexA-lk-PAmCherry with the membrane during the ciprofloxacin-induced SOS response.*** *Quantification from dual-colour SMLM imaging of LexA-lk-PAmCherry and Potomac Red labelled membranes. Histograms show the distribution of the fraction of LexA-lk-PAmCherry localizations within the membrane region per cell for 0.2x, 1x, 2x, and 5x* $MIC^{CIP}$ *(top to bottom) at the indicated time points. The 0 min condition represents unstressed cells for comparison. Mean ± SD is indicated for each condition; n denotes the number of cells analyzed.*

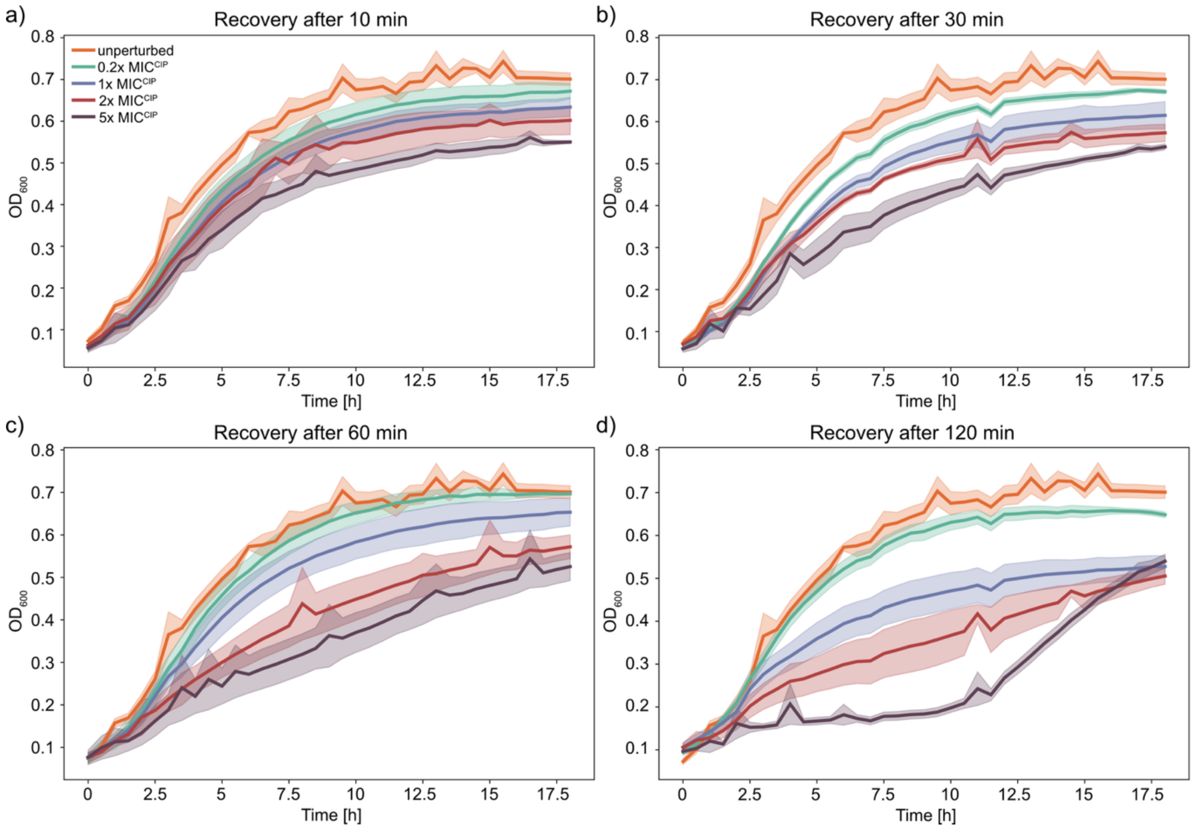

***Supplementary Figure 9 Recovery of E. coli MG1655 following ciprofloxacin stress.*** *Growth curves obtained after exposure to 0.2x, 1x, 2x, or 5x MIC$^{CIP}$ for (a) 10, (b) 30, (c) 60, or (d) 120 min. Following treatment, cells were washed and resuspended in fresh M9 medium, and growth was monitored during recovery. Lines represent the mean of three replicates and shaded areas indicate the SD.*

***Supplementary Table 2 Growth rates during recovery following ciprofloxacin treatment.*** *Growth rates were determined from the initial recovery phase (0–5 h) after ciprofloxacin treatment. Statistical significance was assessed using two-sided Dunnett's test with multiple comparisons correction, with each treatment condition compared against the unstressed control. Significance levels are indicated as ****, p < 0.0001; ***, p < 0.001; **, p < 0.01; *, p < 0.05; and ns, not significant.*

| **timepoint** | **$MIC^{CIP}$** | **Mean growth rate µ [1/h]** | **SD** | **significance** |
|---|---|---|---|---|
| 0 min | 0 | 0.378 | 0.013 | - |
| 10 min | 0.2 | 0.410 | 0.012 | ns |
| 10 min | 1 | 0.401 | 0.013 | ns |
| 10 min | 2 | 0.369 | 0.017 | ns |
| 10 min | 5 | 0.367 | 0.025 | ns |
| 30 min | 0.2 | 0.400 | 0.006 | ns |
| 30 min | 1 | 0.383 | 0.016 | ns |
| 30 min | 2 | 0.331 | 0.018 | ** |
| 30 min | 5 | 0.318 | 0.020 | *** |
| 60 min | 0.2 | 0.376 | 0.022 | ns |
| 60 min | 1 | 0.342 | 0.006 | ns |
| 60 min | 2 | 0.278 | 0.027 | **** |
| 60 min | 5 | 0.246 | 0.012 | **** |
| 120 min | 0.2 | 0.336 | 0.003 | * |
| 120 min | 1 | 0.256 | 0.005 | **** |
| 120 min | 2 | 0.206 | 0.015 | **** |
| 120 min | 5 | 0.125 | 0.011 | **** |

***Supplementary Table 3 Survival assay following ciprofloxacin treatment.*** *Mean CFU/mL for each ciprofloxacin concentration and time point are shorn. N denotes the number of biological replicates, and SEM is given for each condition. Statistical significance was assessed using Dunnett's test, with each treatment condition compared with the unstressed control within the corresponding time-point group. Significance levels are indicated as ****, p < 0.0001; ***, p < 0.001; **, p < 0.01; *, p < 0.05; and ns, not significant.*

| timepoint | $MIC^{CIP}$ | Mean CFU/ml | SEM | n | significance |
|---|---|---|---|---|---|
| 0 min | 0 | 7.52e+07 | 5.20e+06 | 4 | - |
| 10 min | 0 | 9.31e+07 | 2.09e+07 | 3 | - |
| 10 min | 0.2 | 3.50e+07 | 1.15e+07 | 4 | * |
| 10 min | 1 | 7.25e+07 | 1.77e+07 | 3 | ns |
| 10 min | 2 | 5.91e+07 | 1.32e+07 | 5 | ns |
| 10 min | 5 | 1.92e+07 | 8.48e+06 | 4 | * |
| 30 min | 0 | 9.64e+07 | 4.50e+07 | 4 | - |
| 30 min | 0.2 | 7.78e+07 | 3.34e+07 | 3 | ns |
| 30 min | 1 | 7.59e+07 | 1.40e+07 | 3 | ns |
| 30 min | 2 | 1.05e+08 | 5.01e+07 | 5 | ns |
| 30 min | 5 | 2.40e+07 | 1.35e+07 | 4 | ns |
| 60 min | 0 | 1.33e+08 | 9.76e+06 | 3 | - |
| 60 min | 0.2 | 1.19e+08 | 9.26e+06 | 4 | ns |
| 60 min | 1 | 4.54e+07 | 1.56e+07 | 3 | ns |
| 60 min | 2 | 6.20e+07 | 4.49e+07 | 5 | ns |
| 60 min | 5 | 3.55e+07 | 1.06e+07 | 6 | ns |
| 120 min | 0 | 1.68e+08 | 3.79e+07 | 4 | - |
| 120 min | 0.2 | 1.05e+08 | 7.17e+06 | 4 | ns |
| 120 min | 1 | 2.67e+07 | 2.28e+07 | 3 | *** |
| 120 min | 2 | 2.01e+07 | 1.50e+07 | 5 | *** |
| 120 min | 5 | 7.89e+05 | 4.15e+05 | 5 | **** |

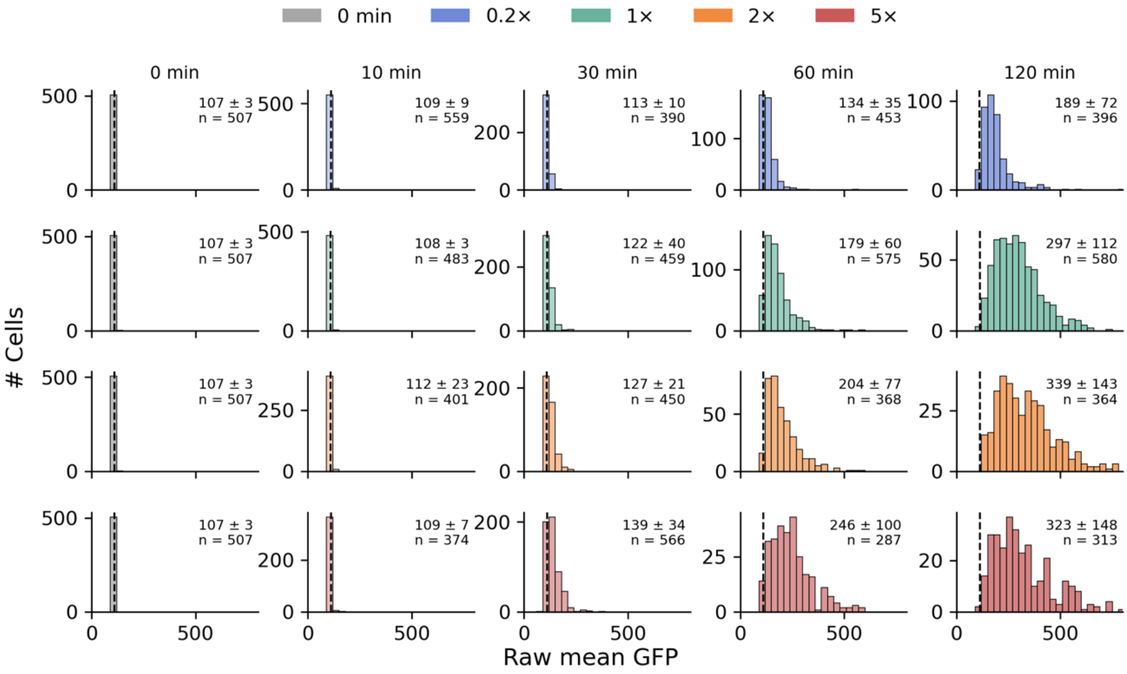

***Supplementary Figure 10 Raw GFP fluorescence signal per cell in E. coli MG1655 sulApΩgfp-mut2 during ciprofloxacin treatment.*** *Histograms show the distribution of raw mean GFP fluorescence per cell following treatment with 0.2x, 1x, 2x, or 5x MIC$^{CIP}$ for the indicated times. Values within each panel indicate mean ± SD and n, the number of analyzed cells. The dashed line indicates the background-derived fluorescence threshold used to classify cells as SOS activated.*

### Supplementary Methods

#### Construction of LexA expression vectors

LexA and LexA-lk-PAmCherry were cloned into pET28a(+) (Novagen), modified to encode a C-terminal octa-histidine (8×His) tag. The corresponding DNA fragments were amplified from pR6K-LexA-lk-PAmCherry-lox71-cm-lox66 using the primer pairs LexA_PAmCherry_inf_F/LexA_His_inf_R and LexA_PAmCherry_inf_F/LexA-PAmCherry_His_inf_R, respectively. The pET28a(+) backbone was linearized using pET28_lin_F and pET28_8xHis_lin_R, which introduced two additional histidine codons into the original hexa-histidine module. PCR products and linearized vectors were verified by agarose gel electrophoresis and purified using the Wizard® SV Gel and PCR Clean-Up System (Promega). Constructs were assembled by In-Fusion HD cloning (Takara) using 50 ng vector and 25 ng insert DNA. Reactions were incubated for 15 min at 50 °C and transformed into chemically competent *E. coli* XL10-Gold cells. Kanamycin-resistant colonies were screened by colony PCR and verified by sequencing using T7 promoter and terminator primers. Confirmed plasmids were transformed into *E. coli* BL21(DE3) for protein expression.

#### Protein expression and purification

*E. coli* BL21(DE3) cells carrying the LexA or LexA-lk-PAmCherry expression plasmid were grown overnight in 50 mL TB medium supplemented with kanamycin. Expression cultures were inoculated in 1 L TB medium to an initial $OD_{600}$ of approximately 0.03 and grown at 37 °C and 120 rpm to an $OD_{600}$ of 0.6–0.8. Protein expression was induced with 1 mM IPTG and the culture was incubated overnight at 18 °C and 120 rpm. Cells were harvested by centrifugation and resuspended in lysis buffer containing 100 mM $Na_2HPO_4$, 200 mM NaCl and 20 mM imidazole, pH 8.0. Cell pellets were stored at −20 °C until purification. After thawing, pellets were resuspended in lysis buffer supplemented with Benzonase® Nuclease and protease inhibitor. Cells were lysed using an Emulsiflex high-pressure homogenizer, and insoluble material was removed by centrifugation for 1 h at 9,782 × g and 4 °C. His-tagged proteins were purified by immobilized metal-affinity chromatography using a 5 ml HisTrap HP Ni Sepharose column. After equilibration with lysis buffer, clarified lysates were loaded at 5 mL $min^{-1}$. Proteins were eluted using a gradient of 0–100% elution buffer containing 100 mM $Na_2HPO_4$, 200 mM NaCl and 1 M imidazole, pH 8.0. Elution fractions were analysed by 12% SDS-PAGE and Coomassie staining. Fractions containing purified protein were dialysed overnight at 4 °C against 100 mM $Na_2HPO_4$ and 200 mM NaCl, pH 8.0, using a 20 kDa MWCO dialysis cassette. Protein concentrations were estimated by comparing Coomassie-stained band intensities with a bovine serum albumin standard series.

**Electrophoretic mobility-shift assay**

Electrophoretic mobility-shift assays were performed with purified LexA and LexA-lk-PAmCherry. Double-stranded DNA substrate containing the LexA boxes of the *recA* promoter were generated by PCR using RecA_SOS_box_F/RecA_SOS_box_R. PCR products were verified on 2% agarose gels and purified using the Wizard® SV Gel and PCR Clean-Up System. Binding reactions contained 0.5 pmol DNA and the indicated protein concentration in 15 µl of binding buffer containing 20 mM Tris-HCl, 0.5 mM EDTA, 200 mM NaCl, 10 mM $MgCl_2$, 5% glycerol and 50 µg $ml^{-1}$ BSA, pH 7.4. Reactions were incubated for 20 min at 25 °C and separated on 5% non-denaturing polyacrylamide gels in 0.5x TBE at 7 mA for 1 h.

Gels were stained with 1x SYBR™ Gold in 0.5x TBE for 30 min and imaged using an Azure Biosystems C300 imaging system with EPI Blue LED excitation. Band intensities were quantified using ImageJ to determine the bound DNA fraction, θ. Binding curves were fitted by nonlinear regression in MATLAB to determine the dissociation constant ($K_D$), maximum binding ($B_{max}$), and Hill coefficient ($n$):

$$\theta = \frac{B_{max}}{1 + (\frac{K_D}{[P]})^n} \qquad \textit{Equation 1}$$

**LexA cleavage assay**

For the RecA-dependent cleavage of LexA and LexA-lk-PAmCherry, 13 µM RecA (NEB) was activated with 1 µM 53 bp oligo (**Supplementary Table 6**) in 20 mM Tris (pH 7.5), 5 mM $MgCl_2$, 1 mM ATPγS and 1 mM DTT and incubated for 2 h on ice. The cleavage reaction was performed in 20 mM Tris (pH 7.5), 5 mM $MgCl_2$, 1 mM ATPγS, 1 mM DTT and 0.5 µM LexA or LexA-lk-PAmCherry after a pre-incubation at 37ºC for 5 min and a subsequent initiation by addition of 1 µM activated RecA filament (RecA*). Cleavage reactions were stopped at indicated time points by addition of 1x SDS sample buffer + DTT and immediate heating to 95°C for 5 min. Samples were loaded on a 4 – 20 % Gradient Precast Tris-Glycine Protein Gels (Bio Rad). Band intensities were analyzed with ImageJ to access the cleaved fraction of LexA and LexA-lk-PAmCherry. The means of three replicates were fitted using an exponential decay function (equation 5) where t represents the time after which the cleavage reaction was stopped and y the fraction of cleaved protein. The decay rate of the protein is described by k. a and c represent the amplitude and offset of protein cleavage, respectively.

$$y(t) = a * e^{-k*t} + c$$ *Equation 2*

***Supplementary Table 4 Plasmids used in this study.***

| **Plasmid name** | **Reference** |
|---|---|
| pR6K-lox71-cm-lox61 | Gift from A. Francis Steward |
| pR6K-sulA-GFPmut2 lox71-cm-lox61 | This study |
| pZA31-sulApΩgfp-mut2 | [1] |
| pSC101-BAD-γβαA | [2] |
| pSC101-BAD-Cre | [2] |

***Supplementary Table 5 Bacterial strains used in this study.***

| **Name** | **Genotype** | **Reference** |
|---|---|---|
| *E. coli* MG1655 | F- λ- rph-1 | Gift from Thorsten Mascher |
| *E. coli* MG1655 lexA-lk-PAmCherry | F- λ- rph-1 *lexA-lk-PAmCherry* | [3,4] |
| *E. coli* MG1655 sulApΩgfp-mut2 | F- λ- rph-1 *Δattλ::sulApΩgfp-mut2* | This study |
| *E. coli* MG1655 lexA-lk-Dendra | F- λ- rph-1 *lexA-lk-Dendra2* | [3] |
| *E. coli* DH5α-pir[+] | *endA1 hsdR17 glnV44 (= supE44) thi-1 recA1 gyrA96 relA1 φ80dlacΔ(lacZ)M15 Δ(lacZYAargF)U169 zdg-232::Tn10 uidA::pir+* | Invitrogen |

***Supplementary Table 6 List of all oligonucleotides. Sequences are shown in 5' to 3' direction.***

| **Name** | **Sequence** |
|---|---|
| sulA_GFP_in_fwd | ACGCTGCCGCAAGCACTCAGGGCGCAAGGGTTATAGGAGAGGCTTTTCATAAAATTCCTTTTAAA |
| sulA_GFP_in_rev | GCTATACGAACGGTACGCTTCCTTTAGCAGGATCCTTATTTGTATAGTTCATCCATGCCA |
| pR6K_lin_fwd | CTGCTAAAGGAAGCGTACCGT |
| pR6K_lin_rev | CCCTTGCGCCCTGAGTG |
| sulA_GFP_rec_fwd | TCACAGGTTGCTCCGGGCTATGAAATAGAAAAATGAATCCGTTGAAGCCTAGGTGAAGTAGGTACCGTTC |

| | |
|---|---|
| sulA_GFP_rec_rev | AAGGTATTAAAAACAACTTTTTGTCTTTTTACCTTCCCGTTTCGCTCAAGATAAAATTCCTTTTAAAATCATAACATAAAAGAATG |
| sulA_GFP_seq_fwd | TGTTCACAGGTTGCTCCG |
| sulA_GFP_seq_rev | GTTAATCACTCTGCCAGATGGC |
| LexA_PAmCherry_inf_F | TTGTTTAACTTTAAGAAGGAGATATACCATGAAAGCGTTAACGGCCAGG |
| LexA_His_inf_R | CTCAGTGGTGATGGTGATGGTGATGGTGGCCGCTGCTGCCCAGCCAGTCGCCGTTGCG |
| LexA-PAmCherry_His_inf_R | CTCAGTGGTGATGGTGATGGTGATGGTGGCCGCTGCTGCCTTTGTACAACTCGTCCATTCCACCTGTAC |
| T7_promoter | TAATACGACTCACTATAGGG |
| T7_terminator | GCTAGTTATTGCTCAGCGG |
| RecA_SOS_box_F | GCCGGGTAATACCGGATAGTC |
| RecA_SOS_box_R | GCTACTGCGTATGCATTGCAG |
| RecA filamentation | TGGCGACGGCAGCGAGGCTTTTTTTTTTTTTTTTTTTTTTTTTTTTTTTTTTT |

### Supplementary Methods References

1. Cui, L. & Bikard, D. Consequences of Cas9 cleavage in the chromosome of Escherichia coli. *Nucleic Acids Res* **44**, 4243–4251 (2016).
2. Wang, J. *et al.* An Improved Recombineering Approach by Adding RecA to λ Red Recombination. *Molecular Biotechnology* **32**, 043–054 (2006).
3. Schärfen, L., Tišma, M., Hartmann, A. & Schlierf, M. Direct Visualization of Four Diffusive LexA States Controlling SOS Response Strength during Antibiotic Treatment. *BioRxiv* (2020).
4. Schärfen, L., Tišma, M. & Schlierf, M. Fast, Simultaneous Tagging and Mutagenesis of Genes on Bacterial Chromosomes. *ACS Synth. Biol.* **9**, 2203–2207 (2020).
